# Action initiation and punishment learning differ from childhood to adolescence while reward learning remains stable

**DOI:** 10.1101/2022.05.05.490578

**Authors:** Ruth Pauli, Inti Brazil, Gregor Kohls, Miriam C. Klein-Flügge, Jack C. Rogers, Dimitris Dikeos, Roberta Dochnal, Graeme Fairchild, Aranzazu Fernández-Rivas, Beate Herpertz-Dahlmann, Amaia Hervas, Kerstin Konrad, Arne Popma, Christina Stadler, Christine M. Freitag, Stephane A. De Brito, Patricia L. Lockwood

**Author notes:** Correspondence should be addressed to: Ruth Pauli, Centre for Human Brain Health, University of Birmingham, United Kingdom., Patricia L. Lockwood, Centre for Human Brain Health, University of Birmingham, United Kingdom.

## Abstract

Theoretical and empirical accounts suggest that adolescence is associated with heightened reward learning and impulsivity. Experimental tasks and computational models that can dissociate reward learning from the tendency to initiate actions impulsively (action initiation bias) are thus critical to characterise the mechanisms that drive developmental differences. However, existing work has rarely quantified both learning ability and action initiation, or it has tested small samples. Here, using computational modelling of a learning task collected from a large sample (*N*=742, 9-18 years, 11 countries), we tested differences in reward and punishment learning and action initiation from childhood to adolescence. Computational modelling revealed that whilst punishment learning rates increased with age, reward learning remained stable. In parallel, action initiation biases decreased with age. Results were similar when considering pubertal stage instead of chronological age. We conclude that heightened reward responsivity in adolescence can reflect differences in action initiation rather than enhanced reward learning.

## Introduction

Adolescence is a time of great change, as young people navigate their way from the dependency of childhood to the independence of adulthood. Theoretical accounts suggest it is a period of risky, impulsive, and reward-seeking behaviour, which is hypothesised to reflect neurobiological changes that lead to heightened reward learning^1–5^. Adolescence is also a high-risk period for the onset of mental disorders^6^, including disruptive behaviour disorders^7^, which are strongly associated with impulsive behaviour and difficulties with reinforcement learning^8^. Internalising problems are likewise associated with difficulties in reinforcement learning^9,10^, and social media use, which can become problematic for some adolescents^11^, and has recently been linked to reward learning mechanisms^12^. However, reward- and punishment-guided behaviour in adolescence is not well understood. This is because distinct psychological processes can manifest in similar overt behaviour, and traditional data analysis techniques are usually not well suited to capturing these covert processes^13^. A myriad of terms have been developed to describe closely related concepts, such as reward learning, risk-taking, and impulsivity^14^, which, though they might reflect similar behaviour, point to distinct psychological processes. Furthermore, these concepts are typically operationalised using questionnaires or summary performance measures from behavioural tasks, which cannot capture the temporally dynamic nature of learning processes^13^. In consequence, our understanding of adolescent behaviour has been impeded by our inability to distinguish between learning processes and other mechanisms that might manifest in similar behaviour, such as response biases. Here, we use computational modelling to distinguish between learning processes (modifying future behaviour based on past experience of reward and punishment) and action initiation or ‘go’ biases (initiating actions impulsively or ‘blindly’, without regard for consequences). We test whether these different mechanisms are separable, and to what extent they exhibit normative developmental differences across late childhood and adolescence in a large and internationally diverse sample.

Computational modelling of learning typically uses reinforcement learning models, which assume that actions and their outcomes become associated through experience, and the learned value of an action then influences the likelihood of repeating that action in the future^15,16^. There has been a relative paucity of computational modelling work focusing on learning in adolescence, and previous studies were not designed to distinguish between learning processes and action initiation biases. Probabilistic learning tasks have suggested an adolescent peak in reward learning^17^ and relatively better reward (versus punishment) learning in adolescents compared to adults^18^. Reversal learning tasks (with changeable outcome probabilities) have pointed to increased punishment learning in adolescents compared to adults^19^, a trough in punishment learning rates in mid-adolescence coupled with a sudden increase in reward learning rates in early adulthood^20^, or peaks in both punishment and reward learning in late adolescence^21^. Together these studies suggest that reward and punishment learning might differ across adolescence, but they provide inconsistent evidence. Part of this variability could be due to different task demands^22^, but it could also reflect the reliance on smaller and non-diverse samples that are not fully representative of adolescents across different countries.

To our knowledge, only one study to date has measured learning in a task design that incorporates requirements both to learn and also to inhibit actions^23^. This study compared reward and punishment learning as well as the tendency to ‘go’ (initiate an action) vs. ‘no-go’ (withhold an action) in children (8-12, n = 20), adolescents (13-17, n = 20), and adults (18-25 years, n = 21). Relative to both children and adults, adolescents exhibited attenuated ‘go’ and Pavlovian (action-consistent-with-valence) biases. Learning was best captured by a generic (not valence-specific) learning rate, and learning rate was not associated with age in this sample. This study suggests that, like learning rates in previous studies, action initiation biases might display developmental differences across adolescence.

In summary, adolescence has been associated with an enhanced ability to learn from reward and possible differences in learning from punishment, but evidence has been inconsistent. The literature is made harder to interpret by small sample sizes, two-group designs (which cannot detect quadratic relationships), and lack of learning contexts designed to assess action biases. Therefore, despite evidence that learning processes can undergo profound changes during adolescence, very little is known about how learning mechanisms differ from action initiation biases during this crucial developmental period or about the robustness of previous findings.

Here, we examined differences in reward and punishment learning and action initiation, using a large, diverse sample (*N* = 742) of youths aged 9-18 years recruited from across Europe. Participants viewed a series of abstract 3D objects and had to learn by trial-and-error whether to respond (‘go’, to win points) or withhold responding (‘no-go’, to avoid losing points) for each object^24,25^ (see Figure 1). We built a set of reinforcement learning models that were fitted to the data using a hierarchical expectation maximisation approach and compared using Bayesian model comparison methods^26–28^. These models varied in terms of whether parameters were included for separate reward and punishment learning rates, action initiation biases (the tendency to respond regardless of expected outcome), and sensitivity to the magnitude (number) of points gained or lost.

**Figure 1.**
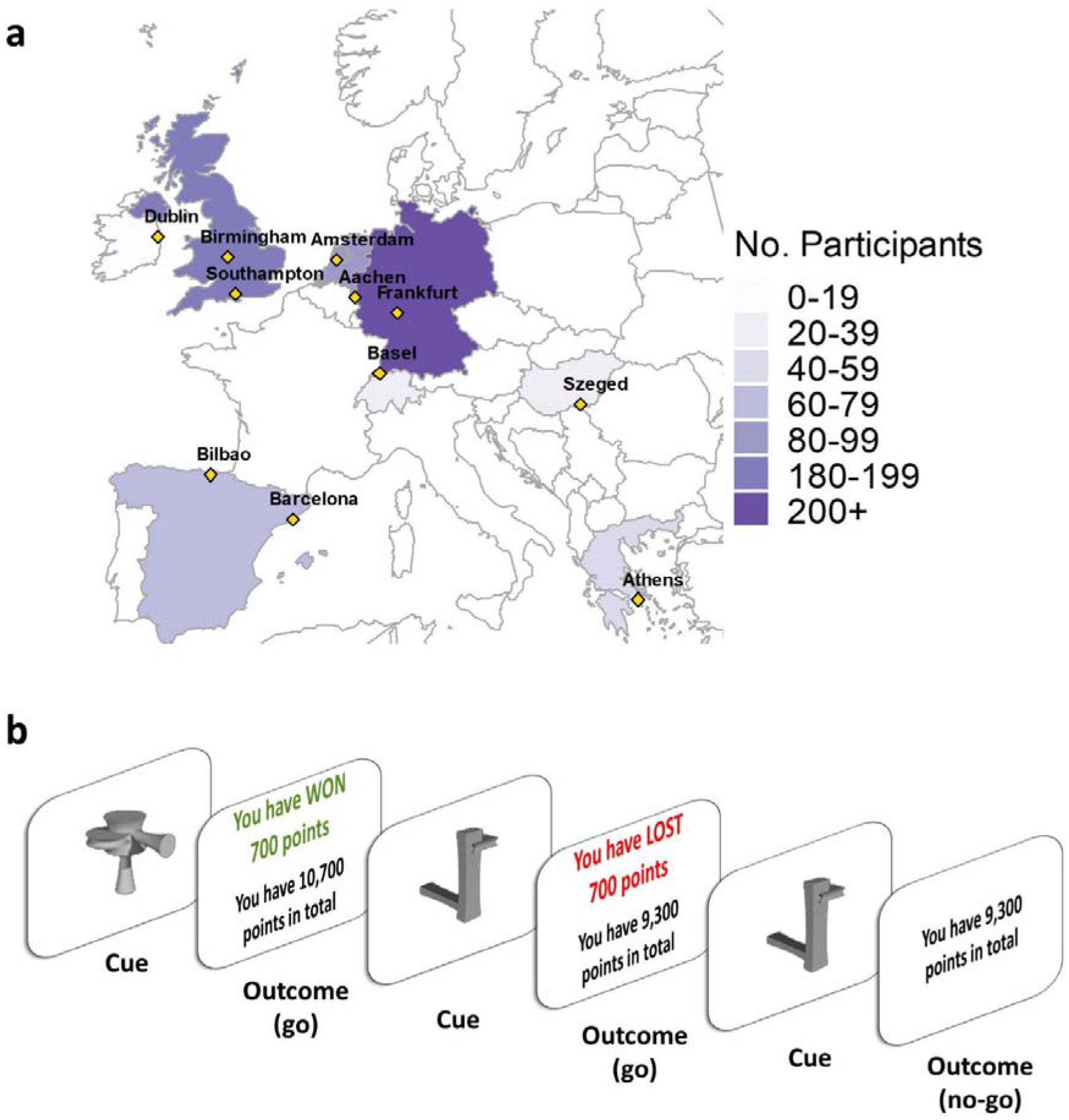
Recruitment sites and learning task. **(a)** Number of participants recruited from each country. Countries are coloured according to the total number of participants, with individual recruitment sites marked in yellow. **(b)** Details of the learning task (shown here in the English language version). The aim of the task was to learn whether to respond or withhold responses to stimuli in order to earn points. Participants learnt by trial and error whether to make or withhold a button press to obtain a reward (points) or avoid punishment (losing points). Eight unfamiliar stimuli were presented individually for 2500ms or until a button press response was made. Responses were followed by feedback on the outcome (1000ms) or a running total alone if the participant did not respond. Each stimulus had a fixed value of +/− 1, 700, 1400, or 2000 points and was shown once per ‘block’ for 10 blocks, with a randomised order within blocks. Thus, four stimuli were associated with reward and four with punishment. Participants started the task with 10,000 points and could theoretically finish with a score between 51,010 and −31,010.

We find that a computational model including separate reward and punishment learning rates, a constant action initiation bias (that measures the tendency to ‘go’ vs. ‘no go’ regardless of reward or punishment on each trial), and a single outcome magnitude sensitivity parameter best explains behaviour. Strikingly, we show an asymmetry in learning differences. While reward learning rates remain stable, punishment learning rates increase from childhood to adolescence. In parallel, despite stable reward learning, action initiation biases decrease with age. All results remain the same when replacing chronological age with pubertal stage. These findings point to normative developmental differences in punishment learning and action initiation. They suggest that theoretical accounts positing heightened responses to reward in adolescence should consider differences in impulsive action initiation rather than reward sensitivity or learning. Such findings are critical for our understanding of learning and decision-making in adolescence as well as how learning and action initiation can go awry in the transition from childhood to adolescence.

## Results

We analysed behaviour of 742 participants (491 girls) aged 9-18 years (mean 13.99, SD = 2.48, median pubertal stage ‘late pubertal’) (see Methods) who completed a reward and punishment learning task (see Figure 1). All participants were free from psychiatric disorders. Pubertal status was measured using the self-report Pubertal Developmental Scale (PDS;^29^ see Methods). After modelling the learning task data, we tested associations between age, pubertal status, and participants’ model parameters, as well as behavioural responses. Age was treated as a continuous variable in all these analyses, although for presentation purposes, we divide age into three discrete bins. To test for quadratic associations between age and model parameters, we tested all models with age^2^ included. We first examined whether there were associations between age or pubertal status and sex or IQ. As there were some associations between these measures (see Supplementary materials), we included sex and IQ as covariates of no interest in all analyses of participants’ behavioural responses and model parameters. For each of these analyses, we ran two models to assess developmental changes: one with participant chronological age and one with pubertal status. Six participants who were included in the computational modelling were removed from subsequent analyses due to missing IQ data.

### Computational modelling shows that a model with separate reward and punishment learning rates and an action initiation bias best explain behaviour

Before fitting and comparing the computational models we analysed participants’ behavioural responses across the task to test whether participants were able to learn. A generalised linear mixed model (GLMM) (predicting correct responses from age, stimulus repetition number, outcome valence, and covariates; see Methods) revealed a significant main effect of stimulus repetition on the number of correct responses made, with performance improving throughout the task (Odds ratio (OR) = 1.19 [1.17, 1.21], *z* = 18.56, *p* < .001). Thus, participants exhibited learning.

Next, we compared a range of computational models of reinforcement learning to characterise participants’ choice behaviour. In particular, we compared models that varied in terms of a single learning rate or separate learning rates for reward and punishment (influence of recent outcomes on future responses), initial or constant action initiation biases (bias to respond versus not respond on the first presentation of an object, or bias to respond versus not respond across all trials, respectively) and sensitivity to the magnitude of reward, punishment or both (sensitivity to points gained or lost). Models were fitted using a hierarchical expectation maximisation approach and compared using Bayesian model comparison methods^26–28,30^. We constructed seven different models using an iterative procedure to appropriately constrain the model space (see Methods for full details):

1. αβ: single learning rate (α) and temperature parameter (β)
2. 2αβ: reward α, punishment α, β
3. αβb*_i* (1): single α, β, initial ‘go’ bias (*b_i*)
4. αβb*_c* (2): single α, β, constant ‘go’ bias (*b_c*)
5. 2αβ*b_i* or 2αβ*b_c*: reward α, punishment α, β, *b_i* or *b_c* (depending on winner from 3. & 4.)
6. 2αβ*b_i*ρ or 2αβ*b_c*ρ: reward α, punishment α, β, *b_i* or *b_c*, magnitude sensitivity (ρ)
7. 2αβ*b_i*2ρ or 2αβ*b_c*2ρ: reward α, punishment α, β, *b_i* or *b_c*, reward ρ, punishment ρ

Models were compared on exceedance probability, Log Model Evidence (LME), and the integrated Bayesian Information Criterion (BIC_int_). We found that Model 6, which included separate learning rates for reward and punishment, a constant action initiation bias, and a single (valence-insensitive) magnitude sensitivity parameter, best explained behaviour (see Figure 2). This model had the highest exceedance probability (0.99) and the highest LME (−34066.81), and performed similarly to model 5 on BIC_int_, which had the lowest absolute BIC_int_. We further validated the winning model using parameter recovery and model identifiability procedures (see Methods for details) and showed good recovery and identifiability for the winning model (see Figure 2 and Supplementary materials). We also examined observed and modelled behavioural performance as predicted by the winning computational model and showed that our model was able to reproduce participant behaviour (see Supplementary materials).

**Figure 2.**
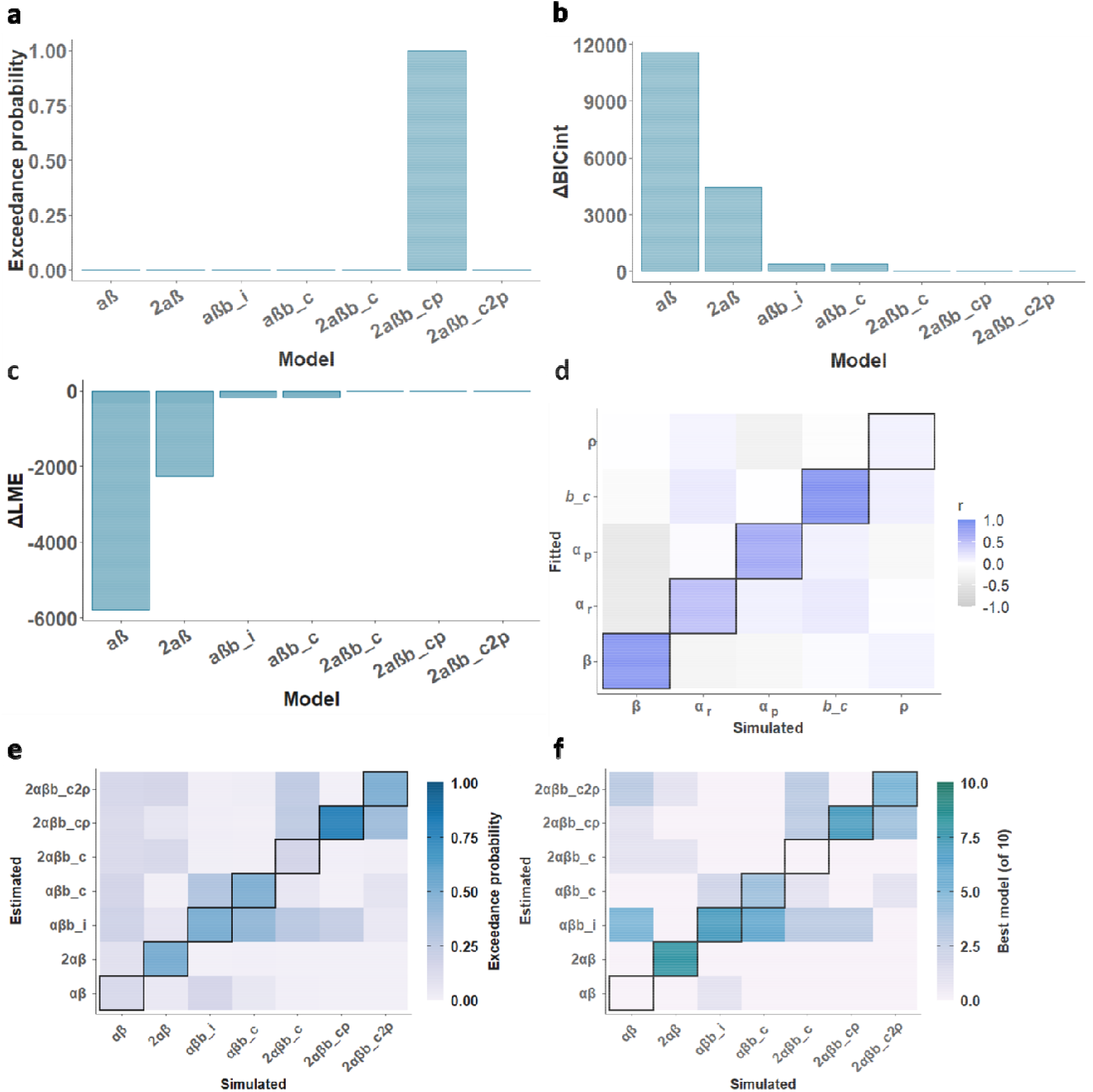
Model performance and validation. **(a)** Exceedance probability for the seven computational models that comprised the model space. The winning model was the 2αβ*b*ρ model, with separate reward and punishment learning rates, a constant action initiation bias, and a magnitude sensitivity parameter. **(b)** ΔBIC_int_, relative to the winning model (2αβ*b_c*). **(c)** ΔLME, relative to the winning model (2αβ*b*ρ). Model 6 (2αβ*b_c*ρ) won on two of the three performance measures (exceedance probability and ΔLME) and performed similarly to model 5, which had the lowest absolute ΔBIC_int_. We therefore selected Model 6 as the winning model **(d)** Parameter recovery. The confusion matrix represents Spearman correlations between simulated and fitted (recovered) parameters. Each parameter exhibited a significant positive correlation between its true and fitted values, with *r* values ranging from 0.1 – 0.83 (shown on the lower diagonal) **(e)** Exceedance probability from the model identifiability procedure. The diagonal represents the probability of each model having the best fit to its own synthetic data. The winning model (2αβ*b_c*) was highly identifiable from other models. **(f)** Number of runs where each model was selected as the best fit for data generated by each model in the model identifiability procedure. The diagonal represents the number of runs each model was selected as the best fit for its own data. The winning model (2αβ*b_c*) was the best fit to its own data.

### Punishment learning rates increase with age, while action initiation biases decline

Next, we assessed whether the parameters from the winning computational model varied as a function of age, using a GLMM predicting correct responses from age, stimulus repetition number, outcome valence, and covariates. Strikingly, age was strongly associated with increased punishment learning rates (*β* = 0.10 [0.05, 0.15], *z* = 4.26, *p* < .001), and lower action initiation biases (*β* = −0.20 [−0.28, −0.12], *z* = −4.78, *p* < .001; see Figure 3). Importantly, reward learning rates did not differ significantly with age (*β* = 0.01 [−0.06, 0.07], *z* = 0.17, *p* = .86). To confirm the strength of these associations, and obtain strength of evidence for any null effects, we calculated Bayes factors using the BIC method^31^ and linear mixed effects regression models, with age removed from the null model. We observed very strong evidence for the associations between age and punishment learning rate (BF_10_ = 336.00, BF_01_ = 0.003) and between age and action initiation bias (BF_10_ = 6545.00, BF_01_ = 0.0002). In contrast, there was no evidence for associations between age and reward learning rate (BF_10_ = 0.07, BF_01_ = 14.30, substantial evidence in support of the null).

**Figure 3.**
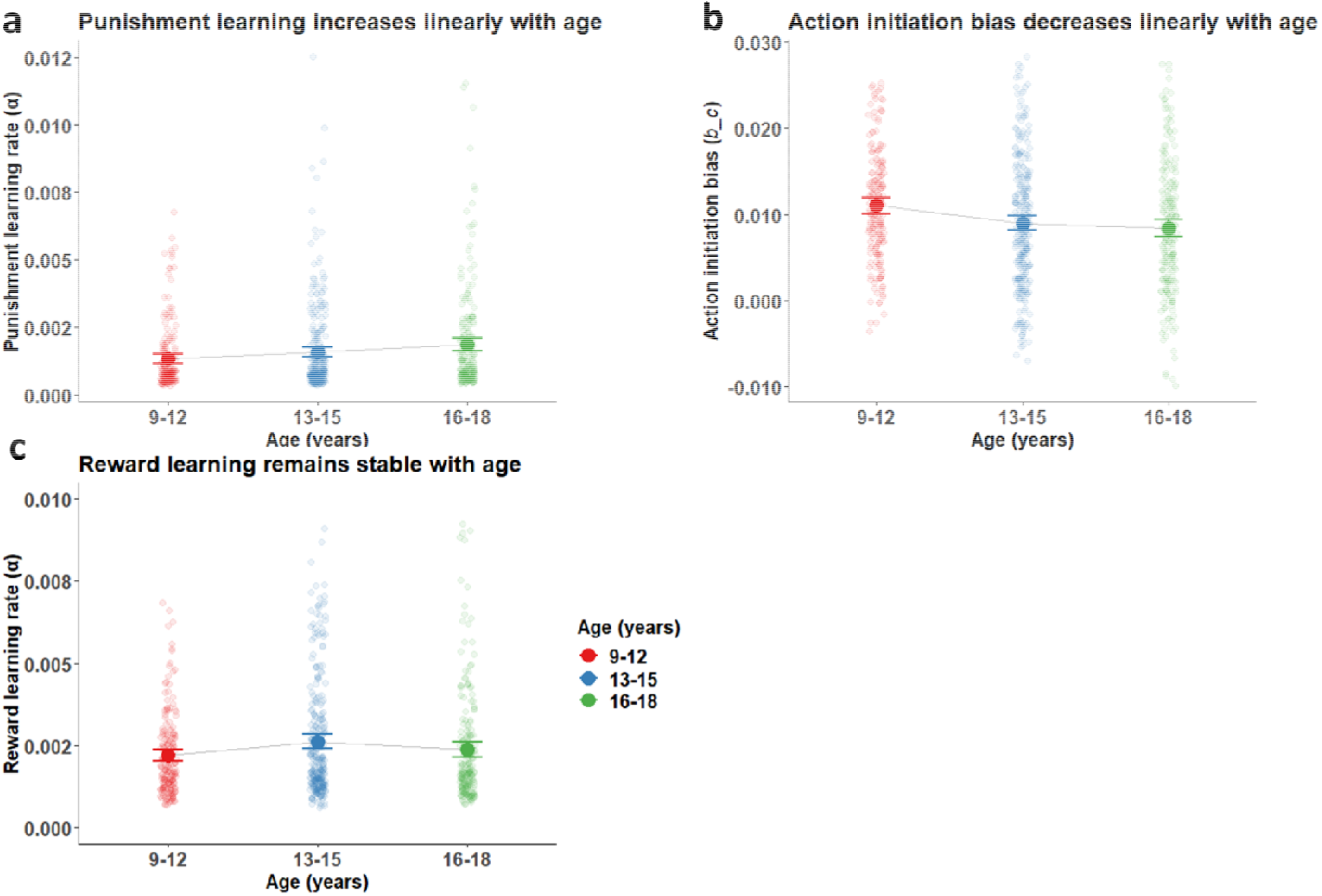
Age differences in action initiation bias and punishment learning, but stable reward learning. **(a)** Punishment learning rate across three age groups. Punishment learning rates increased linearly with age (*β* = 0.1 [0.05, 0.15], *z* = 4.26, *p* < .001, BF_10_ = 336.00, BF_01_ = 0.003). **(b)** Action initiation bias across three age groups. Action initiation biases declined linearly with age (*β* = −0.20 [−0.28, −0.12], *z* = −4.78, *p* < .001, BF_10_ = 6545.00, BF_01_ = 0.0002). **(c)** Reward learning rates across three age groups. Reward learning rates remained stable with age (*β* = 0.01 [−0.06, 0.07], z = 0.17, *p* = .86, BF_10_ = 0.07, BF_01_ = 14.30), including no significant quadratic effect (*β* = -0.83 [−2.37, 0.70], *z* = −1.07, *p* = .29). Points and errors bars represent means and 95% confidence intervals of the means for each group, with raw data represented by smaller points. Division into age groups is for presentation purposes only; age was treated as a continuous variable in all analyses.

We observed a weaker negative relationship between magnitude sensitivity and age (*β* = −0.09 [−0.17, −0.01], *z* = −2.26, *p* = .02), and no relationship between age and temperature parameter (*β* = 0.002 [−0.07, 0.08], *z* = 0.06, *p* = 0.95). Bayes factors showed anecdotal evidence in support of the null for magnitude sensitivity (BF_10_ = 0.51, BF_01_ = 1.97) and substantial evidence for no difference in the temperature parameter (BF_10_ = 0.04, BF_1_ = 26.20).

Lack of associations between age and model parameters might also reflect non-linear associations, especially for reward learning (See Figure 3c). We therefore tested for quadratic effects of age by adding age^2^ terms to the models. However, none of the model parameters exhibited significant quadratic associations with age (temperature parameter: *β* = 0.99 [−0.87, 2.85], *z* = 1.04, *p* = .30). Reward learning rate: *β* = -0.83 [−2.37, 0.70], *z* = −1.07, *p* = .29). Punishment learning rate: *β* = 0.37 [−0.80, 1.54], *z* = 0.62, *p* = .54). Action initiation bias: *β* = −0.57 [−2.63, 1.49], z = −0.54, *p* = 0.59). Magnitude sensitivity: *β* = −0.10 [−2.10, 1.90], *z* = −0.10, *p* = .92).

Next, we assessed whether pubertal stage also predicted differences in punishment learning and action initiation (Figure 4). We re-ran the same models with pubertal stage rather than chronological age. These analyses revealed a similar positive association with punishment learning rate (*β* = 1.41×10^−4^ [6.20×10^−5^, 2.20×10^−4^], *z* = 3.48, *p* <.001), a negative association with action initiation bias (*β* = −0.001 [2.00×10^−3^, 5.00×10^−4^], *z* = −3.40, *p* = .001), and no significant association with reward learning rate (*β* = 5.4×10^−5^ [−5×10^−5^, 1.6×10^−4^], *z* = 1.02, *p* = .31). There was also a negative association with magnitude sensitivity (*β* = −0.01 [−0.01, −0.001], *z* = −2.21, *p* = .03) and no significant association with temperature parameter (*β* = −2×10^−6^ [−5×10^−6^ 10×10^−6^], *z* = −1.29, *p* = .20).

**Figure 4.**
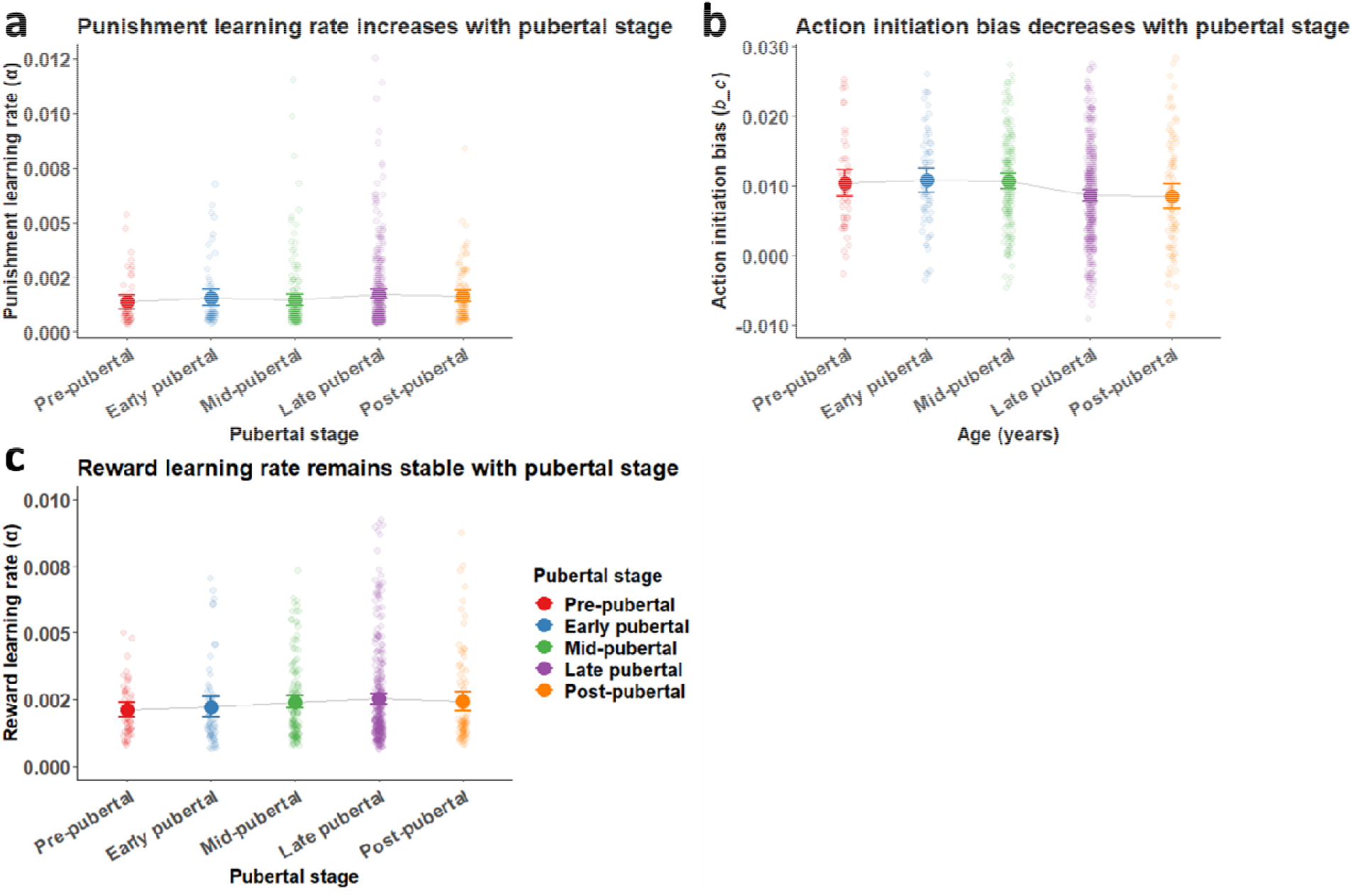
Pubertal maturity differences in action initiation bias and punishment learning, but stable reward learning. **(a)** Punishment learning rates across five pubertal stages. Punishment learning rates increased with pubertal stage (*β* = 1.41×10^−4^ [6.20×10^−5^, 2.20×10^−4^], *z* = 3.48, *p* <.001). **(b)** Action initiation bias across five pubertal stages. Action initiation biases decreased with pubertal stage (*β* = −0.001 [2.00×10^−3^, 5.00×10^−4^], z = −3.40, *p* = .001). **(c)** Reward learning rates across five pubertal stages. Reward learning rates were stable across puberty (*β* = 5.4×10^−5^ [−5×10^−5^, 1.6×10^−4^], z = 1.02, *p* = .31). Points and errors bars represent means and 95% confidence intervals of the means for each group, with raw data represented by smaller points.

### Model parameters predict task performance

We next assessed whether differences in model parameters across age were associated with task performance. Overall task performance (proportion of correct responses) was positively correlated with reward learning rate (Spearman’s r_(693)_ = 0.40 [0.33, 0.46], p <.001) and punishment learning rate (Spearman’s r_(693)_ = 0.67 [0.63, 0.71], p <.001) and negatively correlated with action initiation bias (Spearman’s r_(693)_ = −0.26 [−0.33, −0.18], p <.001). Temperature parameter values and magnitude sensitivity were also negatively correlated with task performance (Spearman’s r_(693)_ = −0.39 [−0.45, −0.32], p <.001, and r_(693)_ = −0.19 [−0.26, −0.12], p <.001, respectively). Correlations between model parameters and reward and punishment task performance are shown in Supplementary materials.

### Behavioural responses confirm age differences in reward and punishment learning

To further confirm that our model accurately captured behaviour, we examined whether age was associated with ‘model-free’ behavioural responses over stimuli repetitions (Figure 5). Age was a significant positive predictor of overall learning (GLMM: age by stimuli repetition interaction: OR = 1.02 [1.01, 1.04], *z* = 2.49, *p* =.01) and older participants also made more correct responses in total (OR = 1.08 [1.04, 1.11], *z* = 4.58, *p* <.001). However, this age-related learning improvement was specific to learning from punishment outcomes (age by repetition by valence interaction: OR = 1.09 [1.05, 1.13], *z* = 4.65, *p* <.001). By contrast, learning from reward outcomes remained stable with age. To quantify the strength of evidence for this stable pattern, we calculated a Bayes factor using the BIC method^31^, by repeating the GLMM model for reward trials only (and removing the valence term), then repeating this reward-only regression with the age*repetition interaction removed. This generated strong support for the stability of reward learning across age (BF_01_ = 57.80; very strong evidence in support of the null).

**Figure 5.**
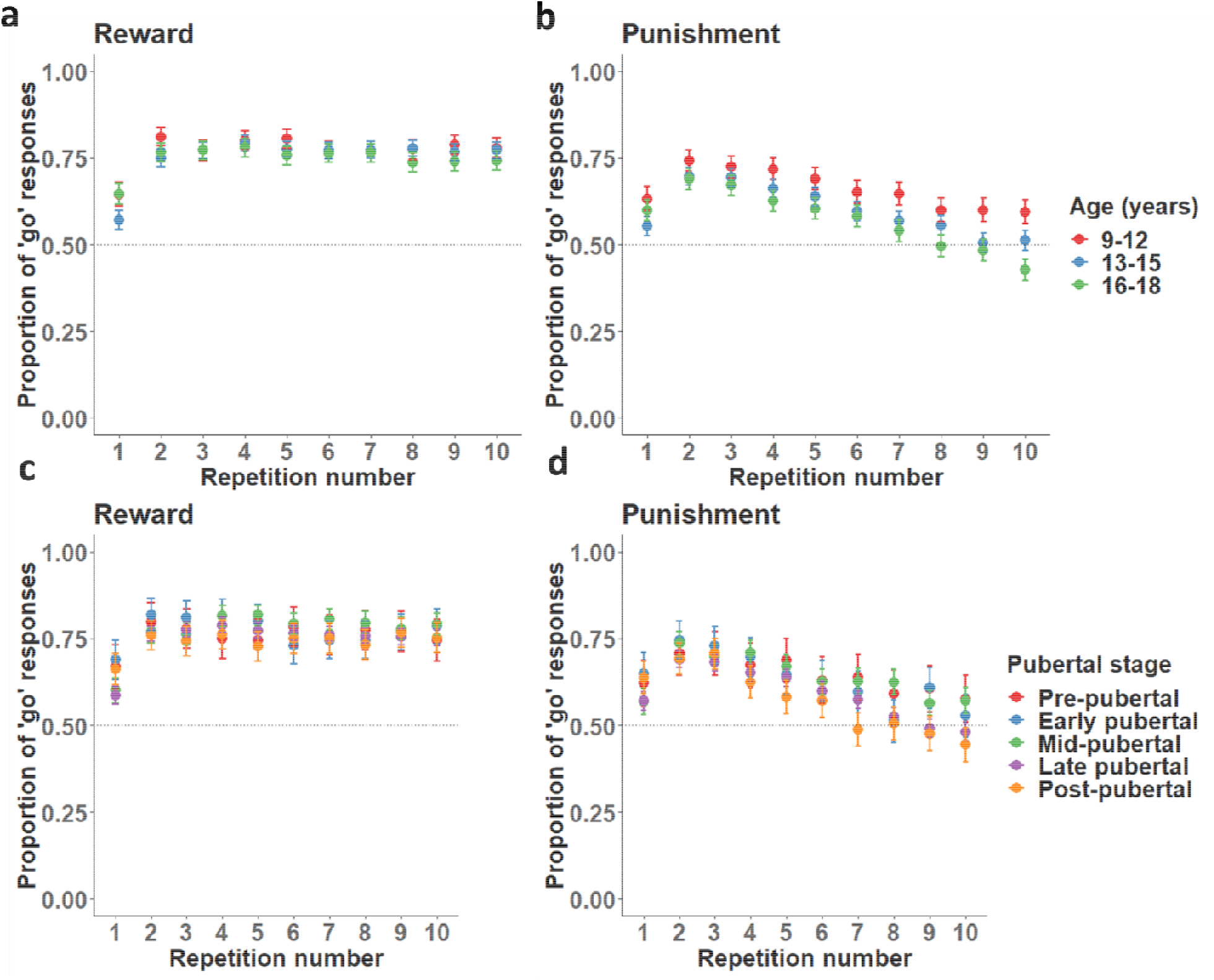
Reward and punishment responding across stimulus repetitions, by age and pubertal stage. **(a)** Proportion of ‘go’ responses to reward stimuli across repeated stimulus presentations, for three age groups. **(b)** Proportion of ‘go’ responses to punishment stimuli across repeated stimulus presentations, for three age groups. **(c)** Proportion of ‘go’ responses to reward stimuli across repetitions, by pubertal stage. **(d)** Proportion of ‘go’ responses to punishment stimuli across repetitions, by pubertal stage. In all panels, points represent means and error bars are 95% confidence intervals of the mean. Dashed lines indicate chance performance. Note that ‘go’ responses are correct for reward and incorrect for punishment stimuli; thus, learning is demonstrated by increasing responses to reward and decreasing responses to punishment stimuli.

In line with our model parameter approach, we tested for quadratic effects of age on behavioural responses. Although this slightly improved the model fit (ΔBIC = −27.79, *p* = .003), the age^2^ term was not a significant predictor of correct responses (OR = 0.99 [0.96, 1.02], *z* = −0.41, *p* = .68) or of overall learning (age^2^ by repetition interaction: OR = 0.99 [0.97, 1.01], *z* = −0.87, *p* = 0.39). However, we did observe a significant age^2^ by repetition by valence interaction (OR = 1.08 [1.04, 1.12], *z* = 3.98, *p* <.001) as well as the significant age by repetition by valence interaction (OR = 1.09 [1.05,1.13], *z* = 4.52, *p* <.001), suggesting that the punishment-specific improvement in learning was partially non-linear.

Since feedback during the task was given in the form of point scores, we also checked for age-related improvements in point score. As expected, older participants gained more points than younger participants overall (robust linear mixed effects regression: *β* = 0.16 [0.09, 0.23], *z* = 4.27, *p* < .001).

### Age-related improvement in punishment learning is not better explained by pubertal development

We next examined whether these age-related improvements in punishment learning were also observed for pubertal stage. Similar to age, pubertal stage was positively associated with overall performance (OR = 1.06 [1.03, 1.10], *z* = 3.71, *p* <.001), and with improved learning (OR = 1.03 [1.01, 1.05], *z* = 2.95, *p* = .003). However, we did not observe a significant pubertal stage by repetition by valence interaction (OR = 1.04 [0.10, 1.07], *z* = 1.87, *p* = .06), suggesting that the punishment-specific improvement in learning was better captured by age than by pubertal stage. Furthermore, the model using age was a better fit to the data than the model using pubertal stage (ΔBIC = −137.22).

## Discussion

Adolescence is often considered as a period of heightened sensitivity to reward^1–5^. Using a large, well-characterised, multi-country sample, we demonstrate that, in fact, reward learning rates remain stable across adolescence whilst the tendency to initiate actions decreases. Moreover, punishment learning rates increase across adolescence, with the oldest adolescents learning the most rapidly from punishment feedback. These findings remained the same when we replaced chronological age with pubertal status, and we found evidence that these differences in model parameters reflected linear associations across adolescence rather than quadratic effects. Together, our findings suggest that the tendency to initiate actions and learn from punishment shifts from late childhood across adolescence and that future research should account for changes in action initiation when evaluating differences in valenced processing of reward and punishment. Our findings also demonstrate these associations robustly by testing a large and geographically diverse sample.

These results highlight the importance of distinguishing between valenced learning mechanisms and action initiation biases. While previous research has demonstrated heightened reward learning in adolescence^17,18^, we demonstrate that apparent reward-oriented behaviour can sometimes reflect action initiation biases, rather than reward learning processes. Knowledge of these developmental differences is an important prerequisite for understanding how adolescent development can go awry, for example in behavioural disorders, where there appear to be disruptions in reinforcement learning^8,32^. It is plausible that adolescent-onset psychopathologies represent aberrant developmental pathways, in which these normative increases in punishment learning and declines in action initiation biases are disrupted. These are important directions for future research.

One consideration is whether action initiation biases are themselves influenced by the prospect of a rewarding outcome, since there are forms of impulsivity that occur specifically in situations where a possible reward is anticipated^33,34^. Since ‘go’ responses in the current study necessarily occur in the context of possible reward, it is possible that the action initiation bias reflects a type of reward-related impulsivity. However, we have two reasons to suspect that this is not the case. First, in contrast to the classic go/no-go paradigm (where ‘go’ responses are required substantially more often than no-go responses), our task used equal numbers of go-for-reward and no-go-for-punishment trials. This means that ‘go’ responses were not particularly associated with reward in this context. Second, we tested a model that captured sensitivity to reward magnitude, but this model was outperformed by a model with a generic magnitude sensitivity. This further suggests that there was no sensitivity to reward driving behaviour other than that captured by the learning rate. These considerations do not support a role for reward in triggering the action initiation bias. Future studies could include ‘go to avoid punishment’ and ‘no-go to gain reward’ conditions to capture the full influence of action biases on reward or punishment responses in a large sample^23^. However, the action initiation bias we observed appears to be a genuine action bias, rather than a deliberate strategy or an indirect effect of reward facilitating action.

Previous research has painted a mixed picture of punishment learning in adolescence, with different studies reporting decreases^20,35^ and increases in punishment learning during the adolescent period^19,21^. It is likely that these differences at least partially reflect variation in task design; in particular, having a higher or lower learning rate can be more or less beneficial depending on the task^22,36^. We observed a positive correlation between accuracy and punishment learning rates across age, suggesting that higher punishment learning rates as seen in older adolescents were more optimal for this task. Thus, the higher punishment learning rates exhibited by older adolescents are indicative of better overall performance. Importantly, however, we did not see increases in reward learning rates across adolescence, although these too were correlated with overall performance. Therefore, the higher punishment learning rates were not simply a reflection of higher general ability on the task, but rather seem to reflect a more specific ability to recall previous punishments and inhibit responses as a result. Crucially, we observe these results in a large and diverse sample of adolescents, providing substantive support for developmental differences in punishment learning.

Although there have been previous reports of heightened reward learning in adolescence^17,18^, the only other study to use a go/no-go design did not observe separate learning rates for reward and punishment^23^. By contrast, our winning model did contain separate learning rates for reward and punishment, demonstrating an asymmetry in learning. However, the lack of an age effect for reward learning in the current study and the lack of a separate learning rate for reward in previous studies^23^ both suggest that reward learning rates are not related to age in a context where action initiation biases can occur. It is theoretically possible that a strong action initiation bias would remove the need for reward learning, since participants could ‘default to go’ and then simply learn from punishment. Again, however, there was a clear association between the reward learning parameter and task performance, and when action initiation biases were lowest in older participants, there was no increase in reward learning rates. This suggests that reward learning was necessary for better performance, even if it did not improve with age. Moreover, for all parameters where we observed differences across development, we saw the same associations when considering pubertal stage. This further suggests that these differences are part of the developmental process, rather than only a reflection of chronological age.

Our study has several strengths. It is among the first to test how action initiation biases and learning differ concurrently across the full spectrum of adolescence, using a learning context that manipulates the requirement for learning and action initiation, something that has often been neglected in computational modelling studies of learning. We used a very large, mixed-sex sample (*N* = 742), which was nationally and linguistically diverse, carefully screened to be typically developing in terms of psychiatric functioning, and well characterised in terms of social background. We built and tested several different plausible models of learning and used multiple measures to validate them. We also used measures of pubertal stage as well as chronological age to further elucidate developmental differences in learning. However, we note some limitations to the study. First, our learning task did not contain ‘no-go to gain reward’ and ‘go to avoid punishment’ conditions, meaning that we were unable to assess Pavlovian action biases^23^. Second, outcomes were deterministic, which has generally not been the case in previous studies (except Master et al., 2020). It is possible that the relationship between learning rates and performance in this context is different from that observed when using the more common probabilistic and reversal learning studies^22,36^.

In summary, we tested developmental differences in learning and action initiation biases in a large, cross-sectional sample of typically developing adolescents aged 9-18 years. Behaviour was best explained by a model with separate learning rates for reward and punishment as well as a constant action initiation bias, and we observed normative developmental differences in these parameters, associated with both chronological age and (to a lesser extent) pubertal stage. Specifically, we observed linear declines in action initiation biases and increases in punishment learning across adolescence, combined with stable levels of reward learning. We conclude that adolescents develop an increasing ability to inhibit actions, learn from negative outcomes, and make more selective behavioural responses as they transition through adolescence and approach adulthood. These findings challenge theoretical and empirical accounts that largely focus on enhanced reward processing and suggest that action biases and punishment learning are crucial processes to understand across adolescence.

## Methods

### Participants

Participants were selected from the FemNAT-CD consortium^38^. All participants included in the present analyses had completed the reinforcement learning task, were 9-18 years old, and were classed as typically developing, with no current psychiatric diagnoses (including autism), learning disability, serious physical illness, or histories of disruptive behavioural disorders or ADHD (see Questionnaire measures below). Eight hundred and thirty-nine participants were eligible for inclusion. We screened the data to exclude participants with poor task performance. Five participants never responded, four responded to every trial, six scored below zero points on the task (indicating deliberate punishment-seeking and reward-avoidance), and 96 responded to fewer than half of the reward trials (i.e., trials where responding was the correct behaviour). The final sample thus consisted of 742 youths (491 girls, 251 boys). These participants were recruited from 11 sites across Europe (Aachen: 139, Frankfurt: 140, Birmingham: 103, Amsterdam: 90, Southampton: 89, Bilbao: 55, Athens: 49, Szeged: 33, Basel: 28, Barcelona: 12, Dublin: 4). For LMM and GLMM (i.e., non-modelling) analyses only, we excluded an additional six participants who were missing IQ data. For the analyses of model parameters and age, we excluded 41 participants with values more than three standard deviations from the mean on one or more model parameters.

All participants provided written informed consent (if over the age of consent in their country) or written informed assent, with written informed consent provided by a parent or guardian. Participants received a small monetary or voucher reimbursement in line with local ethical approvals^39^. This payment was not linked to task performance.

### Questionnaire and interview measures

Participants were assessed for current and past psychiatric and behavioural disorders using the K-SADS-PL clinical interview^40^ (see Supplementary materials). Participants were only eligible for the current study if they were assessed as typically developing according to the K-SADS-PL. IQ was assessed with the vocabulary and matrix reasoning subscales of the Wechsler Abbreviated Scale of Intelligence^41^ at English-speaking sites, or with the vocabulary, block design, and matrix reasoning subscales of the Wechsler Scale for Children (participants <17 years) or Wechsler Adult Intelligence Scale (17-18 years;^42^).

Pubertal stage was assessed using the self-report Pubertal Developmental Scale (PDS;^29^), which assesses growth of body and facial hair, change of voice, and menstruation. Each item is rated on a scale from 1 (*not yet* started) to 4 (*seems* complete). These subscales are then summed to yield an overall pubertal stage score: pre-pubertal (1), early pubertal (2), mid-pubertal (3), late pubertal (4) or post-pubertal (5).

Socioeconomic status (SES) was assessed based on parental income, education, and occupation. Assessments were based on the International Standard Classification of Occupations (International Labour Organization; www.ilo.org/public/english/bureau/stat/isco/) and the International Classification of Education (UNESCO; uis.unesco.org/en/topic/international-standard-classification-education-isced). Human ratings and computer-based ratings were combined into a factor score using principal component analysis. A clear one-dimensional structure underlying the different measures could be corroborated using confirmatory factor analysis (comparative fit index = 0.995; root mean square error of approximation = 0.035). Reliability of the composite SES score was acceptable (Cronbach’s α = 0.74). To account for economic variation between countries, the final SES score was scaled and mean-centred within each country, providing a measure of relative SES. Missing data were imputed by statisticians at the Institute of Medical Biometry and Statistics (Freiburg, Germany), as described in Supplementary materials.

### Learning task

Participants completed a ‘passive avoidance’ reinforcement learning task on a computer in a quiet testing room. The task was adapted from two previous studies^43,44^ and presented in E-Prime^45^. The aim of the task was to gain points by pressing a button when presented with ‘good’ objects (to earn points) and withholding responses when presented with ‘bad’ objects (to avoid losing points). In order to maximise their point score, participants thus had to learn through trial-and-error which objects were associated with reward and which with punishment. There were eight different objects in total, four associated with rewards and four with punishment, with values of +/−1, +/−700, +/−1400, or +/−2000 points. The point value associated with each object was fixed and did not change throughout the task. The eight objects were each presented 10 times in a random order (thus 80 trials in total). Each response was followed by feedback on the number of points gained or lost plus the running total; when participants did not respond, the value of the object was not revealed (see Figure 1). Stimuli were displayed for 3000ms or until the participant responded, and feedback (or the running total alone) was then displayed for 1000ms. Participants started the task with 10,000 points and could theoretically obtain final scores between 51,010 and −31,010, although the maximum score obtainable through learning (rather than ‘lucky guesses’) was 46,909. Since scores below zero could only be obtained by systematically responding to punishment instead of reward, participants with scores below 0 points were excluded (see Participants above).

### Model fitting and comparison procedure

Seven different reinforcement learning models were constructed. For each model, rewards were coded as 1, neutral outcomes (when no response was made) as 0, and punishments as −1. First, we constructed a basic reinforcement learning model, in which learning was captured by a single learning rate (α) parameter and a temperature parameter *β*, which captures noisiness in responding. In this model, the expected value *V* of a response on trial *t* is updated with a reward prediction error *PE* scaled by the learning rate α, where the prediction error is the discrepancy between the outcome *r* (1, 0, or −1) and the expected value:

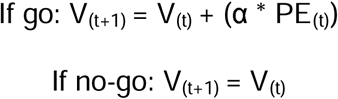

where

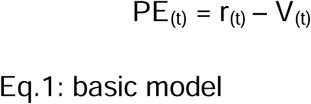

The expected values are then converted to response probabilities using the Softmax equation, where the temperature parameter β adds noise:

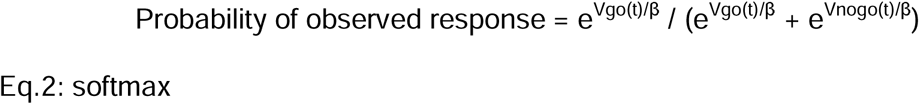

Using the model comparison procedure illustrated in Figure 6, we constructed six further models with combinations of additional parameters. These parameters were separate learning rates for reward versus punishment outcomes (*Eq. 3*), two versions of an action initiation bias towards responding regardless of anticipated outcome (*Eq.4-5*), and one or two magnitude sensitivity parameters, which accounted for sensitivity to the actual point value obtained (*Eq. 6-7*).

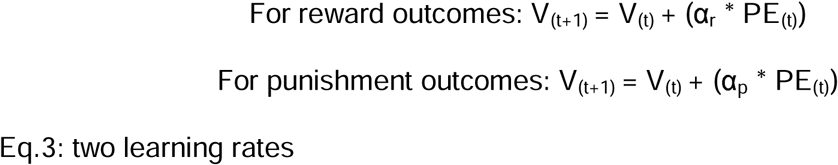

**Figure 6.**
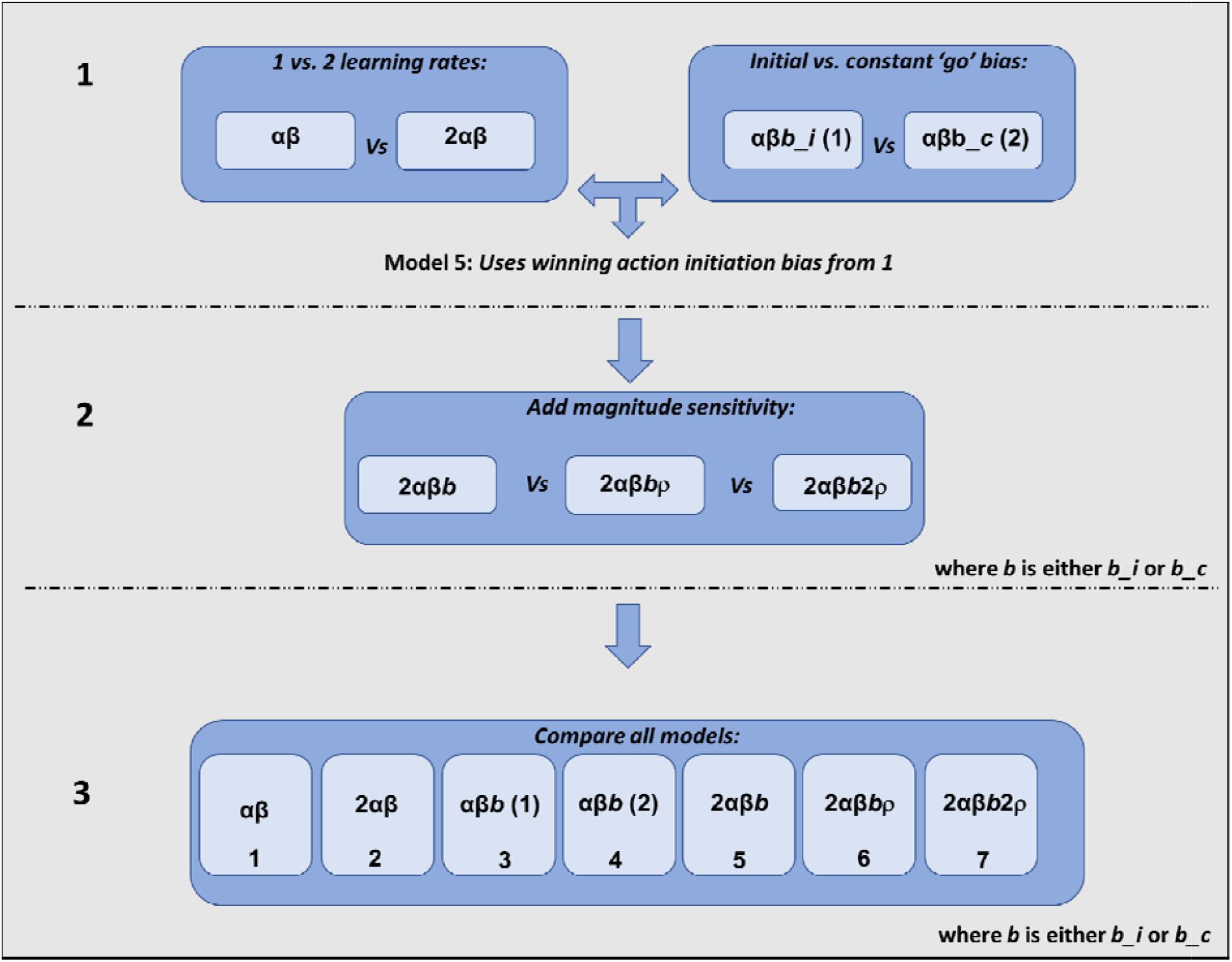
Steps in model construction procedure. In the first step (**1**), models with one versus two learning rates were compared, and separately, models with an initial versus constant action initiation bias were compared. A fifth model was then constructed by combining all parameters from the winning models in step **1** (i.e., one versus two learning rates and the winning action initiation bias). In step **2**, we tested whether model 5 was improved by adding a single magnitude sensitivity parameter (model 6) or separate magnitude sensitivity parameters for reward versus punishment outcomes (model 7). Finally, to confirm that the winning model from step **2** was the best overall model, we compared models 1-7 directly in step **3**.

For models that included the initial ‘go’ bias, the starting value of responding to each object was increased (or decreased) by an amount *b* on the first presentation of the object only:

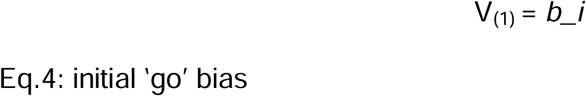

For models that included the constant ‘go’ bias, the value of responding to each object was increased (or decreased) by an amount on each presentation of the object:

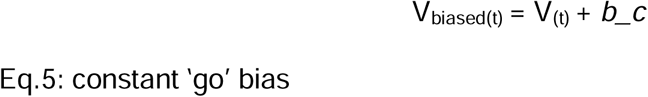

*V*_*biased*_ was used only to calculate the response probability for the current trial, so that the bias did not accumulate over repeated presentations of the object.

For models that included a single magnitude sensitivity parameter, the absolute point score obtained on each trial (re-scaled to be between 0 - 1) was multiplied by a magnitude sensitivity parameter ρ and added to the outcome (which was itself still coded as 1, 0, or −1):

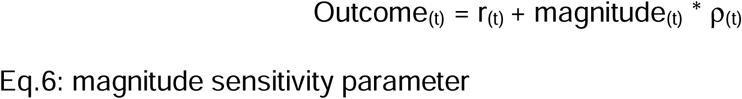

Finally, models that included two magnitude sensitivity parameters applied different magnitude sensitivities to reward and punishment outcomes:

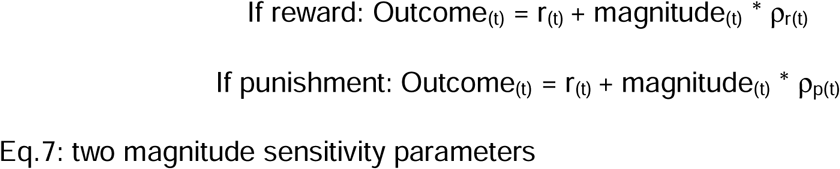

Model fitting and comparison were conducted in MATLAB 2019b (TheMathWorksInc). We used an iterative maximum a posteriori (MAP) approach for all model fitting, in line with previous work using reinforcement learning models^26–28,30^. First, we initialised Gaussian distributions as uninformative priors with a mean of 0.1 (plus noise) and variance of 100. Next, during the expectation step, we estimated the model parameters for each participant using maximum likelihood estimation (MLE), calculating the log-likelihood of the participants’ set of responses given the model being fitted. We then computed the maximum posterior probability estimate, given the participants’ responses and the prior probability from the Gaussian distribution, and recomputed the Gaussian distribution over parameters during the maximisation step. These alternating expectation and maximisation steps were repeated iteratively until convergence of the posterior likelihood, or for a maximum of 800 iterations. Bounded free parameters were transformed from the Gaussian space into native model space using link functions (e.g., a sigmoid function for learning rates).

To compare models, we used Laplace approximation of log model evidence (more positive values indicating better fit^47^) in a random-effects analysis using spm_bms^48^ from SPM8 (www.fil.ion.ucl.ac.uk/spm/software/spm8/). This calculates the exceedance probability, i.e., the posterior probability that each model is the most likely. An exceedance probability over 0.95 provides strong evidence for the best-fitting model. We also calculated the integrated BIC score (BIC_int_) for each model, which penalises more complex models. Lower BIC_int_ scores indicate better performance. MATLAB code for models and model fitting and comparison procedures is available at https://osf.io/d2zp4/.

### Parameter recovery and model identifiability

We used a parameter recovery procedure to ensure that the parameters from the winning model were dissociable from each other, and a model identifiability procedure to ensure that the reinforcement learning models were dissociable from each other^26^. For the parameter recovery procedure, we simulated participant response data only for the winning model, using a range of parameter values between the minimum and maximum possible values for that parameter. Data were simulated for 243 synthetic participants. The winning model was then fitted again to its simulated data using the MAP procedure, and correlations between the parameters used to simulate the data and the recovered parameters (estimated from the simulated data) were checked for correspondence. For the model identifiability procedure, we simulated participant response data for each model in turn, using a range of parameter values within the observed range from the real data. For each of these models, the full set of seven models was then fitted to the simulated data from that model, using the MAP procedure, and this was repeated 10 times. We then created confusion matrices for mean exceedance probability and for the number of times each model won, to check that for each model and its simulated data, the winning model was the one that had been used to generate the data. This procedure confirms that each model is reliably associated with a different pattern of responses from the competing models.

We also generated synthetic behavioural responses using our winning model and its mean parameter values, to check that the real and simulated responses were broadly similar. Finally, as an additional test of the validity of our winning model, we conducted correlations between task performance (number of overall correct responses and correct responses for reward and punishment separately) and each model parameter (Spearman’s correlations, R’s correlation package cor_test function).

### Statistical analysis

All statistical analyses were conducted in R (v. 4.1.1 and v. 4.1.2) through RStudio. First, we investigated associations between age or pubertal stage and the model parameters from the winning model. Since parameter values were not normally distributed, we used robust linear mixed effects regression models using the rlmer function in R. We tested whether each parameter was predicted by age, with IQ and sex as covariates (fixed effects) and varying intercepts for different sites of data collection (random effects). We then checked for quadratic associations with age by adding an age^2^ term to each model. Discrete variables were recoded so that contrasts summed to zero, and continuous variables were z-scored.

To confirm these learning effects matched participants’ behavioural responses, we next used nested linear mixed effects models to assess whether age was related to participants’ changing responses to reward and punishment stimuli over the course of the task. These analyses were conducted using R’s lme4 package glmer function^49^. Participants’ responses were coded as 1 (active response) or 0 (no response) and were predicted from age, sex (0 = male, 1 = female), object repetition number (1-10), and object valence (0 = reward, 1 = punishment) (fixed effects), with varying intercepts allowed for responses grouped by participant nested within site (random effects). All continuous variables were z-scored, and discrete variables (participant response, sex) were recoded so that the two levels summed to zero (e.g., 0 and 1 becomes −0.5 and 0.5). The same analysis was then repeated for pubertal stage, using PDS score as the dependent variable instead of age. In all analyses, IQ and sex were included as covariates. The strength of null effects was interpreted using Bayes factors calculated with the BIC method^31^ and the language suggested by Jeffreys^50^.

## Acknowledgments

R.P was supported by an ESRC post-doctoral fellowship award (ES/V011324/1). P.L was supported by a Medical Research Council Fellowship (MR/P014097/1), a Christ Church Junior Research Fellowship, a Christ Church Research Centre Grant, a Jacobs Foundation Research Fellowship, and a Sir Henry Dale Fellowship jointly funded by the Wellcome Trust and the Royal Society (Grant Number 223264/Z/21/Z). S.A.B.D was supported by an ESRC grant (ES/V003526/1). The FemNAT-CD consortium was funded by the European Commission under the 7th Framework Health Program, Grant Agreement no. 602407. We are grateful to all our participants and their families, and to Jo Cutler, Anthony Gabay, Tobias Hauser, Marco Wittmann, and Stefano Palminteri for helpful discussions and advice.

## Author contributions

R.P conducted all analyses and wrote the manuscript. R.P previously collected data for this study as part of the FemNat-CD consortium. G.K adapted the learning task for use in this study. G.K and I.B collated the learning task data and conducted preliminary data pre-processing and quality checks. M. K-F assisted with computational modelling. J.R contributed substantially to data collection. P.L assisted with coding for the computational modelling analyses, contributed to manuscript preparation, and provided guidance and oversight on all aspects of the analyses. All other authors contributed substantially to study design and/or data collection as part of the FemNat-CD consortium. All authors read and approved the final manuscript.

## Ethics Declarations

The FemNAT-CD project received ethical approval from the relevant local ethics committees, as follows: Aachen: Ethik Kommission Medizinische Fakultät der Rheinisch Westfälischen Technischen Hochschule Aachen (EK027/14). Amsterdam: Medisch Etische Toetsingscommissie (2014.188). Athens: Election Committee of the First Department of Psychiatry, Eginition University Hospital (641/9.11.2015). Barcelona: Child and Adolescent Mental Health—University Hospital Mutua Terrassa (acta 12/13). Basel: Ethik Kommission Nordwest-und Zentralschweiz (EKNZ 336/13). Bilbao: Hospital del Basurto. Birmingham and Southampton: University Ethics Committee and National Health Service Research Ethics Committee (NRES Committee West Midlands, Edgbaston; REC reference 3/WM/0483). Dublin: SJH/AMNCH Research Ethics Committee (2014/04/Chairman (3)). Frankfurt: Ethik Kommission Medizinische Fakultät Goethe Universität Frankfurt am Main (445/13). Szeged (Hungary): Egészségügyi Tudományos Tanács Humán Reprodukciós Bizottság (CSR/039/00392–3/2014). This study was conducted in accordance with the ethical standards of the 1964 Declaration of Helsinki and its later amendments.

