## Supplementary Information for "Action initiation and punishment learning differ from childhood to adolescence while reward learning remains stable"

Correspondence should be addressed to:

**This PDF file includes:**

Supplementary results

Supplementary methods

Figures S1 to S2

Tables S1 to S2

### Supplementary Results

#### Winning computational model tracks learning from reward and punishment

To further confirm the accuracy of our model we generated additional synthetic response data for 125 ‘participants’ using the winning model and its mean parameter values. The simulated responses fell within a similar range to the real responses and, especially on punishment trials, followed a broadly similar trajectory (Figure S1). Thus, the learning rates observed in the winning model are compatible with the behavioural evidence of learning we observed in the participants’ response patterns.

#### Correlations between model parameters and task performance

Correlations between model parameters and task performance (proportion of correct responses) for reward and punishment stimuli are shown in Table S2. All parameters exhibited at least one significant correlation, indicating that all our model parameters meaningfully captured elements of performance.

### Supplementary Methods

#### Parameter recovery and model identifiability procedures

Parameter recovery was conducted on synthetic data generated using the winning model (2αβ*b*ρ). The confusion matrix in Figure 2d shows correlations between the simulated and fitted (recovered) parameter values. All parameters were recoverable, as indicated by significant positive correlations (*p* < .05) between true and fitted parameter values on the matrix diagonal. With the exception of magnitude sensitivity (*r* = 0.11), all parameters exhibited *r* values above 0.50. Model identifiability was conducted similarly, by generating synthetic data from each model and then re-fitting the synthetic data to all models, to determine if the generating model could be identified as the best fit to its own data (Figure 2e-f). This procedure was repeated 10 times per model. The winning model had an exceedance probability of 0.80 for being the best fitting model to its own synthetic data, and was identified as the best model for its own data on 7/10 repetitions.

#### Details of K-SADS-PL clinical interview

The K-SADS-PL^1^ is a semi-structured diagnostic interview used to assess psychopathology in children and adolescents. We conducted interviews separately with participants and with their parents, or another responsible adult informant if a parent was not available. All researchers administering the interview had been trained in its use. After completing the two interviews (parent and participant), we then generated combined parent and child summary ratings of all symptoms (past, present, and lifetime). Where assessors gave discrepant ratings for a symptom, they discussed all available information until an agreement was reached for the summary rating. Except for CD, ODD, and ADHD, where DSM-5 criteria were used, all diagnoses were generated based on the DSM-IV-TR diagnostic criteria, which were current at the outset of the project^2^. For the current study, participants were excluded if they met the diagnostic criteria for any current disorder, or (due to broader consortium requirements) any history of externalising behavioural disorders.

#### Imputation of Missing Data

Missing data were imputed by statisticians at the Institute of Medical Biometry and Statistics (IMBI), a member of the FemNAT-CD consortium. Missing data for the PDS were imputed separately, before the decision was made to impute missing values for other measures. The procedure for the PDS imputation is thus described separately from the other measures. The following description is a standard text provided by IMBI, for use in all FemNAT-CD consortium publications.

Missing values of the PDS score were imputed based on the whole FemNAT-CD sample. It has been shown that missing data in a multi-item instrument is best handled by imputation at the item level^3^. Thus, missing values of the single items were imputed first, and the scores were calculated based on the imputed items. The imputation was done in SAS® version 9.4 using the procedure PROC MI. Imputation by fully conditional specification (FCS) is used, which offers a flexible method to specify the multivariate imputation model for arbitrary missing patterns including both categorical and continuous variables^4^. As the items are measured at an ordinal level, the logistic regression method is specified in the FCS statement. For imputation diagnostics, distribution of the observed and imputed items and scores were checked. The imputation of the PDS items was done separately in males and in females because of sex specific items: item 2 (females and males) and items 4, 5a of the form for females or items 4, 5 of the form for males were imputed respectively. The following variables were included in the imputation model: sex specific items of the PDS as mentioned above and the two remaining PDS items (items 1 and 3), age at PDS and age at informed consent, to impute age at PDS if missing, weight, case/control status, site, and migration status.

Imputation for the remaining measures was conducted separately, following the same procedure as above. The following variables were included in the imputation model: all items of the respective questionnaire, age, IQ, group (case/control), sex (male/female), site, comorbidities (post-traumatic stress disorder (PTSD), attention-deficit/hyperactivity disorder (ADHD), oppositional defiant disorder (ODD), depression, anxiety), and items of other questionnaires if correlated with at least one of the items with ≥.4. For imputation diagnostics, distribution of the observed and imputed items and scores were checked. Data were not imputed for the learning task itself.

### Supplementary Figures

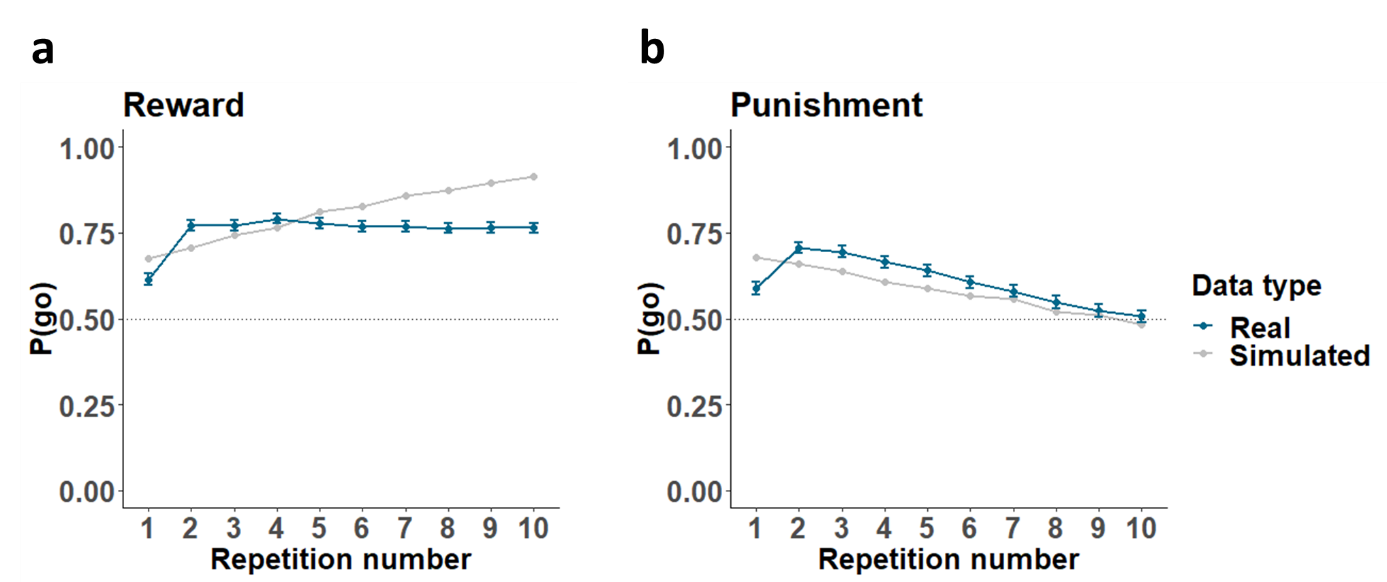

**Figure S1.** **Simulated probability of ‘go’ response to reward and punishment stimuli across 10 stimulus repetitions.** **(a)** Simulated and real probability of ‘go’ responses on reward trials, with simulated data generated using the winning model and its mean parameter values. **(b)** Simulated and real probability of ‘go’ responses on punishment trials. Simulated behaviour broadly mimics real behaviour in both cases.

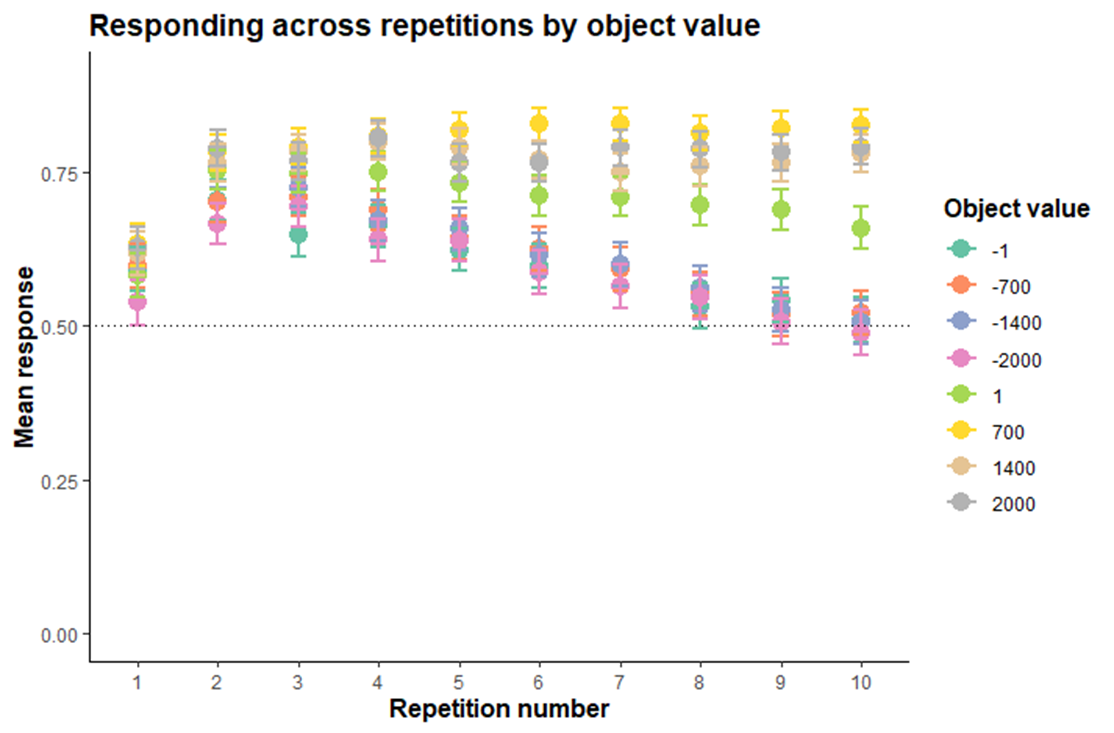

**Figure S2. Responding across repetitions by object value.** Responses to stimuli across repetitions, according to object point value. Responses to reward objects (with values of 1, 700, 1400, or 2000) remain fairly similar after the second presentation, with the exception of the lowest value reward object (1), where responses decrease again towards the end of the task. Responses to all punishment objects decrease steadily after the second presentation.

### Supplementary Tables

**Table S1.** Correlations between demographic variables. Figures are Pearson’s r [95% confidence intervals]

|  | Age | Pubertal status | Sex | IQ |
| --- | --- | --- | --- | --- |
| Age | - | 0.75 *  [0.72, 0.78] | 0.05 *  [−0.02, 0.12] | −0.26 *  [−0.33, −0.2] |
| Pubertal status |  | - | 0.36 *  [0.29, 0.42] | −0.23 *  [−0.29, 0.12] |
| Sex |  |  | - | 0.08 *  [0.00, 0.15] |
| IQ |  |  |  | - |

Notes: * indicates p < .05

**Table S2.** Correlations between model parameters and task performance (proportion of correct responses) for reward and punishment stimuli separately. Figures are Spearman’s r [95% confidence intervals]

|  | β | α_r_ | α_p_ | *b_c* | *ρ* |
| --- | --- | --- | --- | --- | --- |
| Reward performance | −0.59 *  [−0.64, −0.54] | 0.20 *  [0.13, 0.28] | −0.01  [−0.09, 0.06] | 0.82 *  [0.79, 0.84] | 0.30 *  [0.23, 0.37] |
| Punishment performance | 0.12 *  [0.03, 0.18] | 0.19 *  [0.12, 0.27] | 0.57 *  [0.51, 0.62] | −0.86 *  [−0.88, −0.84] | −0.40 *  [−0.46, −0.33] |

Notes: β: temperature parameter, α_r_: reward learning rate, α_p_: punishment learning rate, *b_c*: constant action initiation bias, ρ: magnitude sensitivity. * indicates p < .05
